# No evidence for sleep-dependent memory generalization in a large online sample

**DOI:** 10.64898/2026.03.03.709421

**Authors:** Tiange Lu, Zhuangning Ji, Alexa Tompary, Eitan Schechtman

## Abstract

Memory generalization allows individuals to extract and apply information from prior experiences to novel situations, supporting flexible learning and efficient decision-making. Theoretical models suggest that sleep should facilitate generalization, yet the literature examining its role in promoting generalization is mixed. We recruited 137 participants via Prolific to complete an image-location memory task over two sessions spaced 12 hours apart. Participants were randomly assigned to the Wake group (learning in the morning) or the Sleep group (learning in the evening). In Session 1, participants learned the location of stimuli on the screen and were tested on their memory five minutes later. Twelve hours later, in Session 2, they were tested on their memory again. Stimuli consisted of 160 images from eight semantic categories and were strategically positioned on-screen to test the effects of generalization on retrieval (i.e., category-based memory distortions and biases). After the delay, retrieval was less accurate and demonstrated more generalization. However, these effects were mostly independent of Group, with some evidence for enhanced generalization following a period of wakefulness over sleep. Generalization was also driven by time of day, with more generalization in the evening relative to the morning. Taken together, our results, based on a large online sample, do not support a role for sleep in promoting memory generalization.

**Significance Statement:** Behavior is often guided by memories of previous experiences. However, for behavior to be adaptive and flexible (e.g., when encountering never-before-seen stimuli), regularities about the world must be extracted from these memories. This process, termed memory generalization, has been hypothesized to rely on sleep. We used a large online sample to test sleep’s role in generalization and found no support for this hypothesis. Our results suggest that sleep and wakefulness contribute to generalization equally, with the latter potentially having a larger contribution.

## Introduction

Memories for past events include information on statistical regularities that are instrumental for future planning and informed decision-making (Schacter et al., 2007). During offline periods that follow encoding, memories are transformed through a process termed memory consolidation. Consolidation on the systems level is a complex process that involves both strengthening of specific memories and extraction of the generalized gist, supported by a transformation of their neural underpinnings (Diekelmann & Born, 2010; Lewis & Durrant, 2011; Paller et al., 2021). The Complementary Learning Systems framework proposes that consolidation involves the cross-talk between two stores of information – a “fast learner” which acquires memories initially, and a “slow learner” which gradually embeds the newly learned information into existing semantic networks (McClelland et al., 1995, 2020). The cross-talk between these two stores is thought to occur during sleep, emphasizing its crucial role in memory generalization (Buzsáki, 1989; Inostroza & Born, 2013; Lewis & Durrant, 2011; Witkowski et al., 2020).

However, empirical support for sleep’s role in generalization has been mixed (see Lerner & Gluck, 2019 for review). Arguably, this is due to multiple operationalizations of generalization across studies, encompassing interconnected yet distinct concepts such as transitive inference (Ellenbogen et al., 2007), statistical learning (Lewis & Durrant, 2011), insight into hidden rules (Lacaux et al., 2021; Wagner et al., 2004), semantic learning (Schapiro et al., 2017), and generation of false memories (Mak et al., 2023; Payne et al., 2009). All these forms of generalization involve the extraction of statistical regularities from multiple exemplars and applying them to future decision-making, although the nature of the regularities and of the decisions vary considerably. Sleep may play a role in some forms of generalization but not others, and its contribution may also vary based on other dimensions, such as memory strength (14, 17).

In this study, we tested the effects of sleep on generalization (operationalized as category-based gist learning) using a large sample and a well-replicated task (Tompary et al., 2023; Tompary & Thompson-Schill, 2021). Previous studies examining the effects of sleep on category-based gist learning have produced mixed results. The Deese– Roediger–McDermott (DRM) paradigm has been extensively used for studying false memories. Participants first learn a set of words (e.g., table, sofa, seat) and are then tested on the presented words, but also on a critical lure that was missing from the original list (e.g., chair). Generalization is quantified as the rate of false memories for the critical lure. Some early studies showed increased generalization following sleep (e.g., Payne et al., 2009), while others showed the opposite effect (e.g., Fenn et al., 2009) or no overall effect (e.g., Diekelmann et al., 2010). A meta-analysis (Newbury & Monaghan, 2019), complemented by a recent pre-registered, well-powered online study (Mak et al., 2023), demonstrated that sleep indeed increased the probability of forming false memories in the DRM task, barring certain boundary conditions. Notably, generalization under this operationalization is expressed through erroneous decisions (misremembering a critical lure), whereas real-life generalization ideally leads to adaptive behavior, with generalization errors being the exception rather than the rule.

Another task that examined category-based gist learning explicitly defined several categories of visual objects (i.e., satellites) that include multiple exemplars that have shared, as well as unique, features (Schapiro et al., 2017; Siefert et al., 2024; Tandoc et al., 2021). Here, too, results have been mixed: initial findings that nocturnal sleep improved memory for shared features (Schapiro et al., 2017) were not replicated when changes to the learning regimen were made (Tandoc et al., 2021). In line with these results, biasing reactivation during sleep prioritized unique features over shared ones, demonstrating a potential detrimental effect of reactivation during sleep on generalization (Siefert et al., 2024).

We aimed to test sleep’s impact on generalization using a large online sample and a task that operationalizes generalization with high validity. This task, which was based on the experiments described in Tompary & Thompson-Schill (Tompary & Thompson-Schill, 2021), reconciles a persistent tension in prior literature on memory generalization between abstract rule learning and real-world semantic memory. Although it does not require an encoding strategy that heavily relies on generalization, it enables participants to leverage category knowledge to improve their performance. Participants (N = 137) learned the on-screen location of exemplars belonging to eight distinct categories (e.g., sea animals; Figure 1a). Most items of each category were clustered together in a specific part of the screen (i.e., consistent objects), while some were positioned outside of that part (i.e., inconsistent objects; Figure 1b). If participants position the inconsistent objects closer to the category cluster center, this would be considered as evidence for generalization. Inconsistent objects were either typical category exemplars (e.g., BIRDS – Cardinal) or atypical exemplars (e.g., BIRDS – Ostrich), with the former expected to show stronger generalization-based errors than the latter. In our version of the task, we also included objects that belonged to the learned categories but were not shown during learning and had no veridical on-screen location. Placing these objects near the category cluster center would also be taken as evidence for generalization. In a mixed design, participants were tested both immediately after learning (Session 1; S1) and again ∼12 hours later (Session 2; S2). This delay occurred either overnight, spanning a standard sleep window (Sleep Group), or during the day, spanning normal waking hours (Wake Group). The participants in the Sleep group were directed to sleep naturally as they usually would, and the participants in the Wake group were directed to stay awake and refrain from napping.

**Figure 1.**
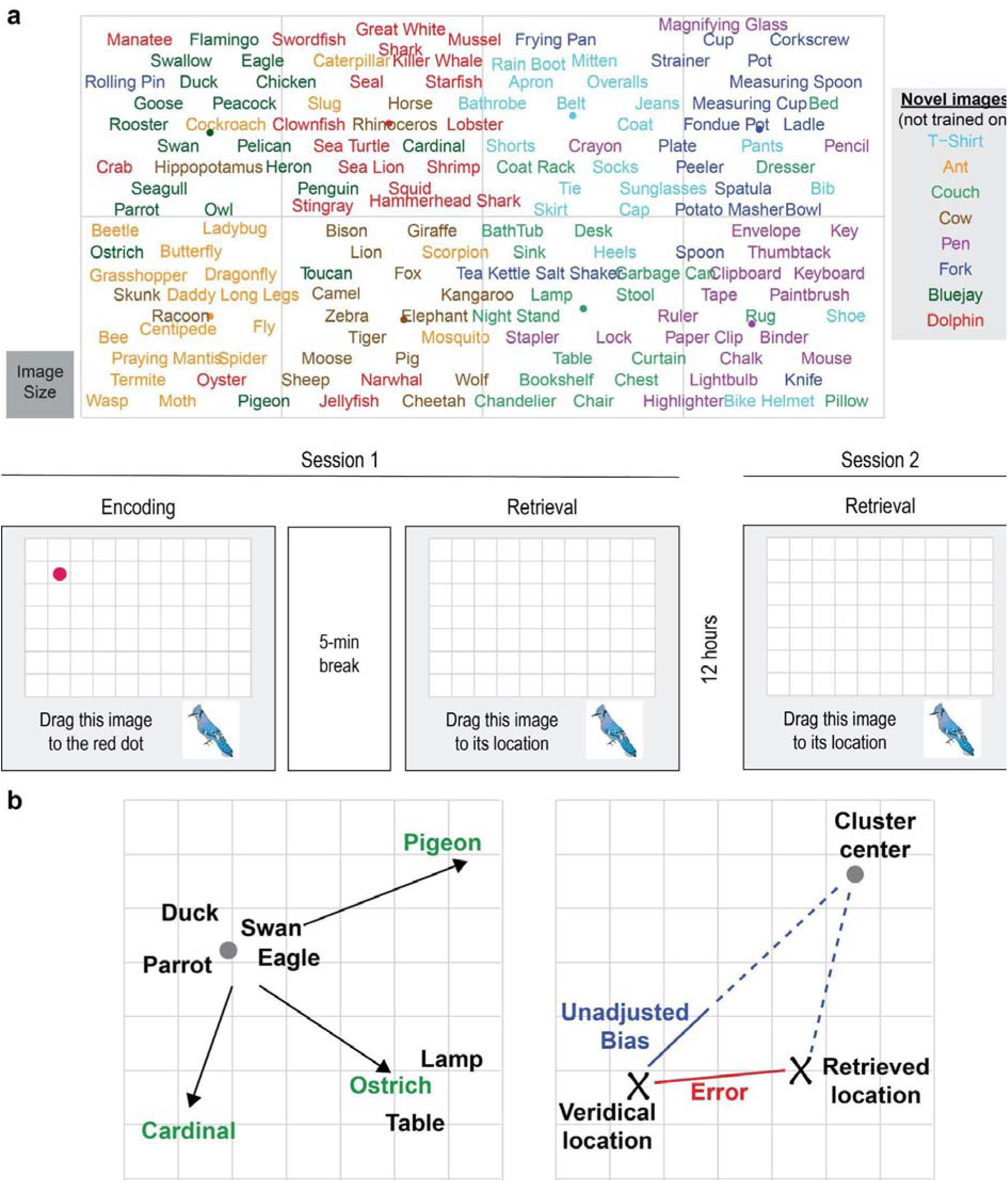
Memory task. **(a)** Images of exemplars belonging to different categories were assigned on-screen locations. The figure shows an example layout for one participant, replacing each image with its label. Images were evenly distributed between eight categories (shown here in different colors for clarity). Image locations were strategically assigned so that most images within a category would be clustered together within a rectangle occupying an eighth of the screen. Participants learned each image’s location over three rounds in Session 1, by dragging the image to a red dot indicating its veridical location. After a 5-minute distraction task, they were tested on their memory. In Session 2, ∼12 hours later, participants retrieved the location of each image again. **(b)** The left panel shows spatial consistency and categorical typicality for an example category (BIRDS). Black font indicates spatially consistent images, and the grey dot shows the category cluster center. Green bolded font indicates spatially inconsistent images, which included typical (e.g., Cardinal) and atypical (e.g., Ostrich) members of the category. The right panel shows the calculation of error and unadjusted bias for an inconsistent image, estimating its bias towards the category cluster center.

Our pre-registered hypotheses (<u>osf.io/qj9ck/</u>) are listed in Table 1. We hypothesized that our results would replicate the main findings of prior research by Tompary & Thompson-Schill (19): In Session 1, regardless of Groups, placement error would be smaller for consistent objects compared to inconsistent ones, demonstrating that category-based information drives performance. Within the inconsistent images, we hypothesized that placement errors would be larger for typical objects compared to atypical ones due to the former being erroneously placed closer to the category cluster center. This would demonstrate that generalization depends on the strength of the relationship between an object and its category. This effect would also be reflected by a measure of bias toward the category center, which would be larger for typical objects compared to atypical ones. We also predicted that there would be a general benefit of sleep for memory, in that the decline in memory performance would be higher for the Wake Group relative to the Sleep Group.

**Table 1.** Pre-registered hypotheses.

| Session 1 Only | Changes between Sessions 1 and 2 |
| --- | --- |
| We will replicate the findings that:<br>1. Consistent Error < Inconsistent Error<br>2. Typical Bias > Atypical Bias<br>3. Typical Error > Atypical Error | The veridical memory for items will generally decline over the delay for both groups, but the Wake group will have a steeper decline compared to the Sleep group. |
| Memory for the locations of images will be biased by prior semantic knowledge (more than chance). | Memory for the locations of images will become more biased by prior semantic knowledge in the Sleep group relative to the Wake group. |
| Positioning of novel typical images, which were not trained on, will be biased by prior semantic knowledge (more than chance). | The difference between typical and atypical bias will increase over the delay, more so for the Sleep group vs. the Wake group. |
|  | Novel typical images, which were not trained on, will become biased toward the cluster center over the delay, more so for the Sleep group relative to the Wake group. |

Our main hypotheses centered on sleep’s effects on generalization. We hypothesized that generalization would increase over the delay, more so for the Sleep Group than the Wake Group. Specifically, we predicted that there would be a greater increase in the placement errors for inconsistent objects in the Sleep Group relative to the Wake Group, demonstrating a general increase in category generalization. This effect would be coupled by a selective increase in bias for the typical (vs. atypical) objects, which would demonstrate that sleep’s effect on generalization is dependent upon the strength of category-item relationships. Finally, we predicted that the change in cluster-centered placement across the delay for the novel objects, which were not learned and had no veridical on-screen location, would be greater for the Sleep Group. Placement of these novel objects close to the category cluster center demonstrates generalization to novel items, for which there is no specific competing memory trace.

## Results

A final sample of 137 participants completed the task, including 67 participants in the Wake group and 70 participants in the Sleep group. The two groups did not differ in the duration of their engagement with the task. The Wake group started Session 1 in the morning (average start time: 9:01 AM ± 38.74 min SD), whereas the Sleep group started Session 1 in the evening (average start time: 9:02 PM ± 34.49 min SD). The duration of Session 1 was 51.82 ± 18.81 min for the Wake group and 51.32 ± 15.97 min for the Sleep group (*t*(133) = 0.17, *p* = 0.87). This session was followed by a delay of 11.66 ± 0.87 h for the Wake group and 12.18 ± 0.84 h for the Sleep group. Despite similar instructions, the delay was significantly longer for the Sleep group (*t*(135) = 3.57, *p* < 0.001). The duration of Session 2 was 14.26 ± 7.47 min for the Wake group and 16.61 ± 6.54 min for the Sleep group (*t*(135) = 1.98, *p* < 0.05).

### Generalization drives memory retrieval of object locations

In Session 1, all participants learned the on-screen locations of 152 images, equally divided between eight categories (Tompary & Thompson-Schill, 2021). Fourteen of the imaged objects within each category were clustered in a rectangular area (Consistent items), whereas five of the images were positioned out of this rectangle (Inconsistent items). Of those five images, two depicted an object highly Typical of the category and three depicted highly Atypical objects. Immediately after learning, participants engaged in a distraction task followed by a memory test. Participants were required to place all images in their correct locations. Objects were placed closer to their veridical position than expected by chance in Session 1 (*t*(136) = 38.79, *p* < 0.001).

We first assessed whether performance on this test differed between groups. Across all images, there were no differences in spatial error between the Sleep and Wake groups in Session 1 (*t*(135) = 0.66, *p* = 0.51). Similarly, errors for Consistent images, Inconsistent images, Typical Inconsistent images, and Atypical Inconsistent images were not different between groups (all *p*s > 0.41). In Session 1, participants across groups remembered the on-screen positions of spatially Consistent images more accurately than spatially Inconsistent ones (*t*(135) = 18.11, *p* < 0.001). This indicates that participants better remembered object locations when these objects were within close proximity of their semantically related counterparts, replicating the effect observed in (Tompary & Thompson-Schill, 2021). Furthermore, the results showed that Inconsistent Typical images exhibit greater bias toward their category cluster center compared to Inconsistent Atypical images (*t*(135) = 7.32, *p* < 0.001). The same effect was reflected by larger errors for Typical images relative to Atypical images (*t*(135) = 5.81, *p* < 0.001). Taken together, these results reflect increased generalization for Typical items in Session 1, as participants’ responses were more strongly biased toward the category center. These results also replicate those observed in (Tompary & Thompson-Schill, 2021). There were no effects of the Group on any of these three effects (all *p*s > 0.08). Taken together, these results demonstrate there were no initial group differences in these metrics. Furthermore, they demonstrate that there are no time-of-day effects (e.g., driven by circadian differences), since Session 1 for the Wake and Sleep groups was scheduled in the morning and evening, respectively.

Approximately 12 hours after Session 1, participants completed a follow-up memory test, which was identical to the first one. Critically, the participants in the Sleep group slept in the interim, whereas the participants in the Wake group remained awake. To test the effects of the 12-h delay and whether sleep vs. wakefulness impacted performance, we ran a series of mixed linear models. First, we replicated the results of (Tompary & Thompson-Schill, 2021) again, this time collapsing over Group and Session. Again, Consistent error was significantly smaller than Inconsistent error (*F*(1,540) = 712.51, *p* < 0.001; Figure 2a), Typical bias was significantly larger than Atypical bias (*F*(1,540) = 73.86, *p* < 0.001; Figure 2b), and Typical error significantly exceeded Atypical error (*F*(1,540) = 47.55, *p* < 0.001).

**Figure 2.**
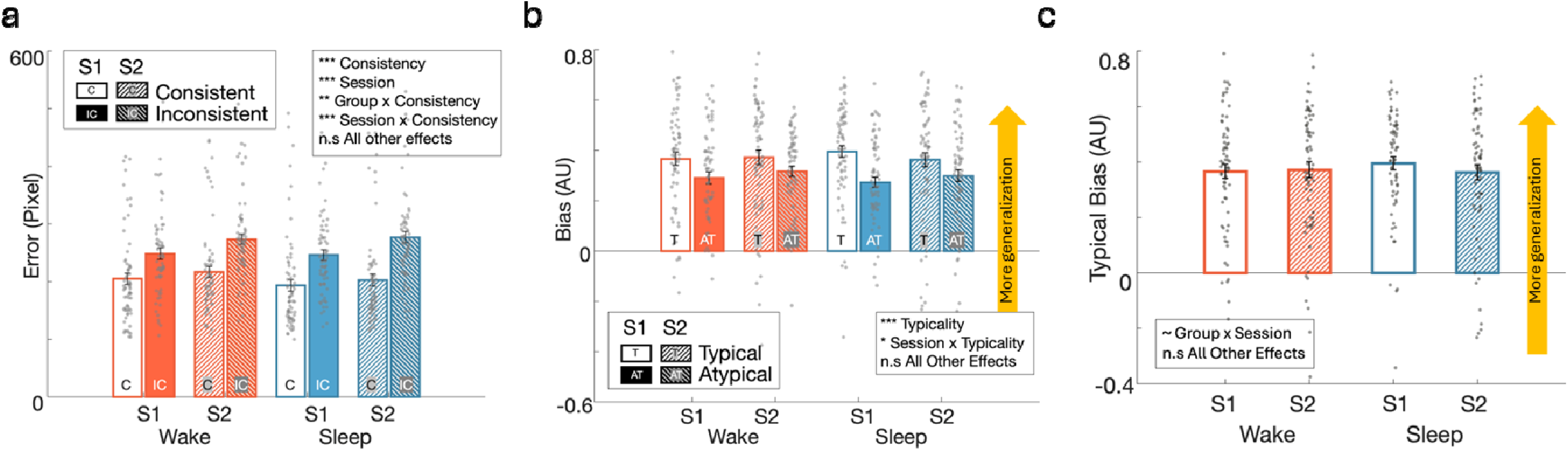
Generalization drives memory retrieval, but is not impacted by sleep. **(a)** Consistent error and Inconsistent error for each Group and Session. Consistent error (lighter bars, marked by C) is significantly lower than Inconsistent error (densely filled bars, marked by IC). Session 2 error is significantly larger than Session 1 error. The significant interaction between Group x Consistency was driven by enhanced Consistency effects for the Sleep group. The significant Session x Consistency interaction was driven by enhanced Consistency effects for Session 2. Critically, there was no three-way interaction between Group, Session, and Consistency. **(b)** Typical bias and Atypical bias for each Group and Session. Typical bias (lighter bars, marked by T) is significantly larger than Atypical bias (densely filled bars, marked by AT). The Session x Typicality interaction was driven by reduced Typicality effects for Session 2. **(c)** Typical bias for each Group and Session. There is a trend toward an interaction of Group and Session, driven by a marginal decrease in bias for the Sleep group following the delay. ∼ *p* < .1, * *p* < .05, ** *p* < .01, *** *p* < .001, n.s non-significant. S1 - Session 1, S2 - Session 2.

Whereas bias was not modulated by the delay (*F*(1,540) = 0.47, *p* = 0.49), error significantly increased between Session 1 and Session 2 (*F*(1,540) = 81.31, *p* < 0.001 in the model including Consistency; *F*(1,540) = 88.39, *p* < 0.001 in the model including Typicality). Similar results were observed when considering different types of errors directly (i.e., using models that included only Group and Session, but did not include Typicality and Consistency as fixed effects). Using these models, we showed that the following types of errors grew larger over the delay: overall error (*F*(1,270) = 37.79, *p* < 0.001), Consistent error (*F*(1,270) = 16.60, *p* < 0.001), Inconsistent error (*F*(1,270) = 92.14, *p* < 0.001), Typical error (*F*(1,270) = 37.59, *p* < 0.001), and Atypical Error (*F*(1,270) = 79.18, *p* < 0.001). Additionally, participants were more likely to place an Inconsistent object closer to the cluster center than to its veridical location in Session 2 relative to Session 1 (*F*(1,270) = 100.3, *p* < 0.001), and the same was true for Typical (vs. Atypical) objects (Typicality x Session interaction effect; *F*(1,540) = 15.9, *p* < 0.001).

### No effects of sleep on generalization

To examine the effects of sleep on memory, we focused on the interactions between Session and Group. We first asked whether sleep improved overall memory (regardless of Consistency and Typicality). If sleep had facilitated memory performance, we would expect a significant interaction effect, with performance improving from Session 1 to Session 2 in the Sleep group more so than in the Wake group. However, this interaction proved non-significant (*F*(1,270) = 0.0056, *p* = 0.94). This indicates that sleep did not improve overall memory relative to wakefulness.

We next tested whether sleep impacted generalization using a similar approach. Generalization was operationalized in several different ways, including the effects of Consistency and Typicality mentioned above. Considering both Consistent and Inconsistent errors separately, there were no interactions between Group and Session (Consistent error, *F*(1,270) = 0.25, *p* = 0.62; Inconsistent error, *F*(1,270) = 0.98, *p* = 0.32). Using a model that includes Consistency, Group, and Session, we found no significant interaction between the three factors, suggesting that sleep did not differentially impact Consistent vs. Inconsistent images, and therefore does not have an effect on generalization (*F*(1,540) = 0.96, *p* = 0.33). However, we did find an interaction of Group × Consistency, indicating that the Sleep group showed larger detriments for Inconsistent vs Consistent items when compared to the Wake group, regardless of Session (*F*(1,540) = 9.40, *p* < 0.01). We also found an interaction of Session × Consistency, indicating that Inconsistent errors increase more than Consistent ones between sessions (*F*(1,540) = 16.92, *p* < 0.001) (see also (Tompary et al., 2020)). The interaction between Group and Session was not significant (*F*(1,540) = 0.14, *p* = 0.71). Examining whether participants placed Inconsistent objects closer to the cluster center relative to its veridical position following the delay, we found no difference between the Wake and the Sleep groups (Session x Group interaction effect; *F*(1,270) = 1.48, *p* = 0.22), and the same was true for Typical (vs. Atypical) objects (Session x Group x Typicality interaction effect; *F*(1,540) = 0.07, *p* = 0.8).

We next considered a different metric of generalization, which is sensitive to the strength of the relationship between an object and its category – Typicality bias. As mentioned above, Typical items were more biased toward the category center than Atypical ones. Testing Atypical bias and Typical bias independently, we considered the effects of Session and Group. For Atypical bias, we found no interaction between the two (*F*(1,270) = 6.54e-5, *p* = 0.99). However, there was a trend toward a Group × Session interaction for Typical Bias (*F*(1,270) = 3.39, *p* = 0.07), with the Sleep group exhibiting a larger decrease in Bias over the delay relative to the Wake group (Figure 2c). Note that this trend is in the opposite direction than expected and suggests that generalization decreased, rather than increased, over sleep. Employing a model including Typicality, Group, and Session, we found no significant interaction between the three (*F*(1,540) = 1.05, *p* = 0.31). There was an interaction between Session and Typicality (*F*(1,540) = 4.52, *p* < 0.05), indicating that bias for Atypical images increased over the delay more than bias for Typical images. In other words, Atypical images were placed closer to the cluster center after the delay (relative to the change for Typical images). The effect of image Typicality (i.e., the difference between Typical and Atypical images) shrank between sessions. All other interaction effects, including the critical interactions between Group and Session, were not significant (*p* > 0.13).

To test the effects of Typicality on error (rather than bias), we employed a model including Typicality, Session, and Group. Here, too, we did not find an interaction between the three (*F*(1,540) < 0.01, *p* = 0.96). In this model, all other interactions were also non-significant (all *p*s > 0.21). Taken together, these results support the notion that sleep had no effect on generalization in this task.

### Some evidence supports a benefit of wakefulness to generalization

Along with Consistency and Typicality, a third metric of generalization was incorporated in our design. Participants were trained on 19 object images for each of the eight groups. A twentieth object from each group – the most typical one (e.g., couch - furniture) – was not included in the Training Session, but was nonetheless tested on both before and after the delay. Since this item has no veridical on-screen location, generalization should lead to its placement near the center of the category cluster.

Unlike other inconsistent category members that show biases towards their category clusters, placement of these items near the cluster center is a purer measure of generalization, because there it is not impacted by interference from the learned on-screen locations. We therefore defined its “error” as the distance from that point. Novel objects were placed closer to their category cluster centers than expected by chance in Session 1 (t(136) = 24.5, p < 0.001).

Using a model incorporating Group and Session, we found that Novel item placement was not modulated by Group (*F*(1,270) = 1.51, *p* = 0.22), but its error decreased between Session 1 and Session 2 (*F*(1,270) = 7.22, *p* < 0.01), indicating generalization. Critically, there was an interaction between Group and Session (*F*(1,270) = 7.99, *p* < 0.01), indicating a larger decrease in error for the Wake relative to the Sleep group. This indicates that the Wake group utilized category knowledge to position Novel Images closer to the category cluster centers more so than the Sleep group after the 12-h delay (Figure 3a).

**Figure 3.**
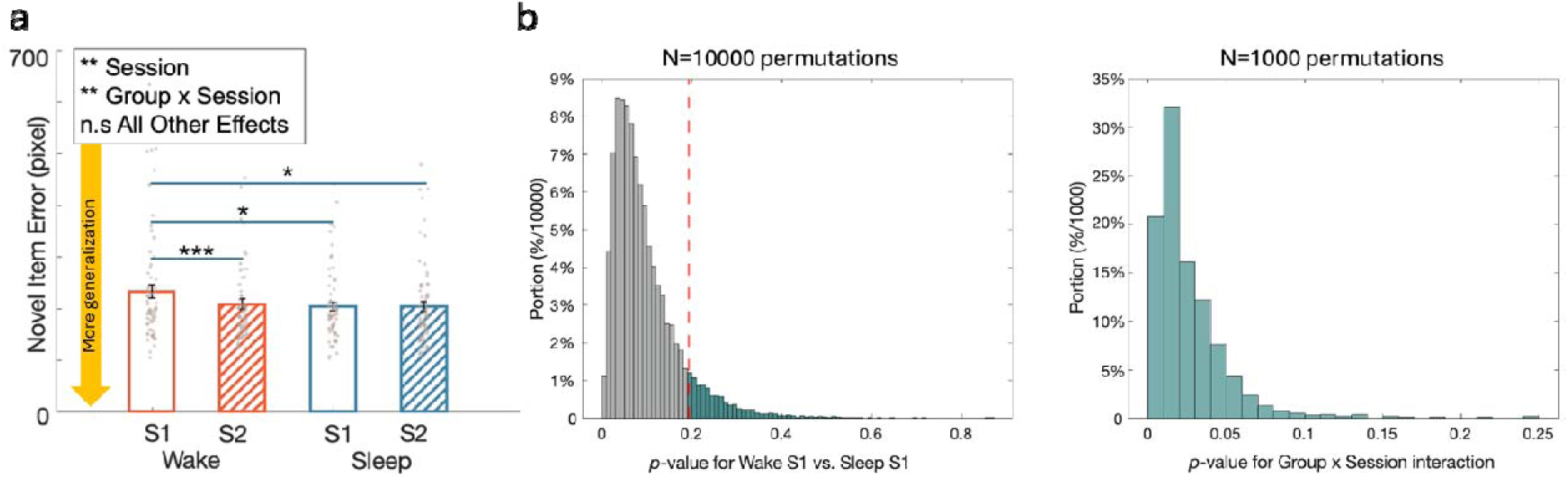
Novel image placement suggests that generalization is enhanced after wakefulness relative to sleep. **(a)** Novel image error for each Group and Session. The significant Session effect and the Group x Session interaction are driven by larger error for Novel Images in Session 1. * *p* < .05, ** *p* < .01, *** *p* < .001, n.s non significant. S1 - Session 1, S2 - Session 2. **(b)** A permutation test for Novel images shows that wakefulness increases generalization (relative to sleep) even when accounting for baseline differences. (left) We subsampled our data over 10,000 permutations to eliminate baseline differences in Novel item error. 10% of permutations with the lowest Session 1 differences (blue bars) were chosen for the subsequent analysis. (right) Within this subset of permutations, the Group x Session interaction remains significant (*p* for 88.8% of permutations < .05).

However, a post-hoc examination of the data revealed differences in Session 1 placement between groups: The Wake Group tended to place the novel object further from the cluster relative to the Sleep group (*t*(135) = 1.93, *p* = 0.06). This difference in baseline scores may have contributed to the reported effect of wakefulness on generalization. To reveal whether these baseline differences could account for the observed interaction, we subsampled the data over multiple permutations. Specifically, we randomly excluded 10 participants from each group across 10,000 iterations, and recalculated the effect of Group on Novel object error in Session 1 for each iteration (Figure 3b, left). Then, we selected 10% of datasets in which Session 1 performance between Groups was most balanced (*p* > 0.19). For these 1,000 subsets, we re-evaluated the Group × Session interaction. The resulting distribution of *p*-values was strongly left-skewed, with 88.80% falling below the 0.05 threshold, indicating that the interaction persisted even after controlling for baseline differences (Figure 3b, right). This suggests that generalization, as reflected by Novel object placement, increases over a period of wakefulness more so than over a period of sleep.

### A trend towards better generalization in the evening

We tested whether the results were driven by the time of day during which tests were taken, which could reflect circadian effects. To do this, we complemented the analysis comparing Session 1 performance across groups with an analysis examining performance at the morning/evening, collapsing over groups. As mentioned above, the Wake group had their Session 1 at the same time range as Session 2 for the Sleep group, and the Wake group had their Session 2 at the same time range as Session 1 for the Sleep group. Collapsing across groups would reveal general time-of-day effects observed across sessions (e.g., effects of fatigue, attention, and motivation). We compared the collapsed data of Wake Session 1 & Sleep Session 2 (morning tests) to the collapsed data of Wake Session 2 & Sleep Session 1 (evening tests), and found generally no time-of-day effect on performance of tasks: no effect on Average error (*F*(1,272) = 0.0028, *p* = 0.96; Figure 4a), Consistent error (*F*(1,272) = 0.15, *p* = 0.70), Inconsistent error (*F*(1,272) = 0.86, *p* = 0.35), Typical error (*F*(1,272) = 0.55, *p* = 0.46), Atypical error (*F*(1,272) = 0.71, *p* = 0.40), Atypical bias (*F*(1,272) = 0.0018, *p* = 0.97). There was a marginal circadian effect for Typical bias (*F*(1,272) = 3.45, *p* = 0.064; Figure 4b), with a larger bias observed in the evening. Similarly, there was a significant difference between morning tests and evening tests in participants’ placement of the Novel images. Participants placed Novel images closer to their category center on evening tests relative to morning tests (*F*(1,272) = 7.29, *p* < 0.01; Figure 4c).

**Figure 4.**
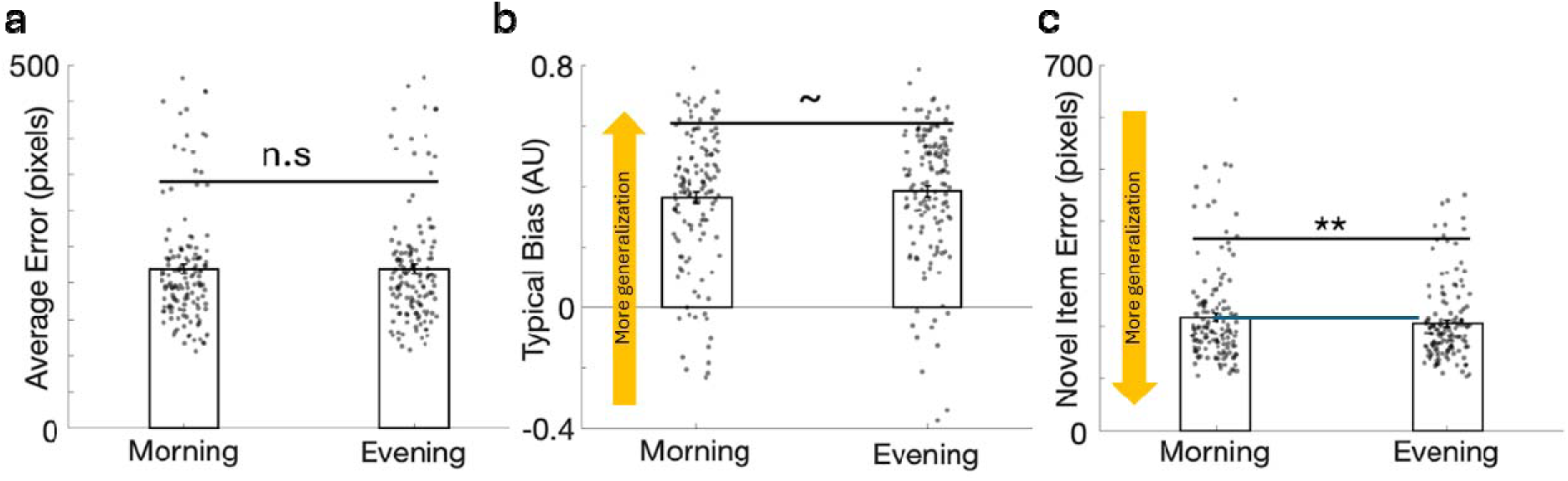
A trend toward more generalization in evening relative to morning tests. **(a)** No significant difference in Average Error between the morning and evening tests. **(b)** There was a trend toward an effect of Typical bias, which was higher in evening tests compared to morning tests. **(c)** Novel Item Error was significantly lower in the evening tests compared to the morning tests, suggesting more generalization in the evenings. ∼ *p* < .1, ** *p* < .01, n.s non significant.

In summary, we replicated key findings from (Tompary & Thompson-Schill, 2021), including greater memory accuracy for spatially consistent over inconsistent images and increased generalization for typical items. No significant Group × Session interactions were found for most measures, suggesting that sleep did not broadly influence performance. However, a significant Group × Session interaction was observed for Novel items, and this effect persisted even after controlling for baseline group differences through permutation testing. Additionally, reliable main effects of Session were observed across both groups, indicating memory deterioration and increased generalization of atypical category members over time, independent of sleep. There was also a marginal trend towards better generalization in the evening sessions compared to the morning sessions, regardless of group.

## Discussion

In this study, we used a well-powered online study to examine the effects of sleep on generalization using a spatial-memory task. Participants learned the on-screen locations of 152 images, which depicted objects from eight categories (19 objects per category). Most images for each category were clustered together, whereas the on-screen location for five items from each category was outside of this cluster. Generalization was operationalized as the tendency to position these inconsistent objects near their category’s cluster center. Participants were tested on all locations immediately, as well as ∼12 hours later. Critically, one group of participants learned in the morning and were tested in the evening (“Wake Group”), whereas the other group learned in the evening and were tested in the morning (“Sleep Group”).

Memory for object positions declined between the two tests, regardless of Group. This decline was not different between the Wake and Sleep groups, suggesting that memory for this task may not be sleep-dependent. When considering the consistent images in each category (those close to the category cluster), sleep did not improve memory relative to wakefulness, indicating a null effect of sleep regardless of generalization. This is surprising, as previous studies using spatial memory tasks did show advantages for sleep relative to a similar amount of time awake (Simon et al., 2022; e.g., Wilhelm et al., 2008). Although we are unsure as to why sleep did not improve memory in this task, it may have to do with the learning regimen, which did not include feedback-guided learning to criterion, but rather a less demanding training routine that involves dragging objects to indicated locations. Although sleep has been suggested to rescue weaker memories (e.g., Drosopoulos et al., 2007; Schechtman, 2024), such as the ones potentially produced by learning without a criterion, others have proposed that it mostly benefits memory in the intermediate range (e.g., Stickgold, 2009). It may, therefore, be that memories were not encoded to a sufficient degree to benefit from sleep in our task.

The critical question addressed in this study is whether sleep promotes memory generalization. We used multiple measures of memory generalization to explore this question. First, we considered the changes in memory placement for inconsistent images (i.e., those that were not positioned near the category cluster center). Error rates for consistent images were lower than for inconsistent images, demonstrating that participants used category-based information to guide their placements. Although both consistent and inconsistent errors increased following the delay, inconsistent errors grew to a larger degree, demonstrating a higher reliance on category regularities at the 12 h mark. Crucially, however, there were no significant differences between the Sleep and Wake groups, suggesting that generalization was not impacted by sleep. Similar null results were obtained when using another measure of generalization – category bias for typical and atypical objects. Of the five inconsistent objects, two were typical category exemplars and three were atypical category exemplars. Positioning of typical objects was biased toward the category cluster more so than for atypical ones. Unlike for inconsistent errors, this bias was not modulated by the delay. Importantly, there were no effects of sleep on this measure of generalization.

A major limitation of these measures is that the study design pits specificity and generalization against each other. Generalization is measured as a deviation from placing an object in its veridical location, i.e., a mistake. Participants in our task were instructed to encode specific image locations. Generalization would have negatively impacted performance, and participants with perfect memories would not have been expected to show any generalization at all: they would remember the veridical locations of each item regardless of whether it is consistent or inconsistent with its category’s statistical regulation and regardless of typicality. These measures of generalization therefore reflect a tug-of-war between item-specific memory and generalization. This issue is methodological in nature and not inherent in the definition of generalization; real-life generalization is best observed in novel situations in which the existing knowledge structures can be applied to improve decision-making. Specific-item memory may not play any part in such applications, and certainly does not conflict with the expression of generalization.

To overcome this issue, we introduced a final measure of generalization. The memory tests included a single typical image for each category that was not shown during the Learning Phase, and therefore has no veridical location. For this memory, there was no competition between generalization and specificity, and positioning it close to the category cluster would not be incorrect (as there is no correct response for such an image). Participants positioned the novel image closer to its category cluster than expected by chance. Furthermore, they placed it closer to the category center after the delay, supporting the notion that generalization increases with delay. Interestingly, the effect of delay on generalization was modulated by Group, with the Wake group showing a larger increase in generalization relative to the Sleep group. Although there were baseline differences in pre-sleep generalization between the groups, subsampling analyses confirmed that this unexpected result is statistically valid. Therefore, our results suggest that wakefulness – rather than sleep – may promote generalization.

Another factor that may drive generalization is the time of day. Studies have shown that generalization tends to occur more in the morning relative to later in the day (e.g., Tandoc et al., 2021; Xie et al., 2018). Our results conflict with these findings. When collapsing over groups and comparing morning and evening tests, we show that despite comparable total error, typical bias was marginally higher in the evening. Additionally, novel objects were placed closer to the category cluster center in evening tests. These time-of-day differences may have limited our ability to detect sleep’s effect on generalization: more generalization in the evening (Session 1 for the sleep group) relative to the following morning session (Session 2) may have masked a benefit of sleep. However, as described above with regard to the analysis involving Novel images, the contribution of wakefulness to generalization holds even when accounting for differences in Session 1 generalization.

One limitation of the present study is the absence of objective sleep recordings, such as polysomnography or actigraphy. Consequently, we cannot verify exact sleep duration, architecture (e.g., slow-wave sleep vs. rapid eye movement sleep), or quality for participants in the Sleep Group, nor can we definitively rule out brief daytime naps in the Wake Group. Because we did not collect objective physiological recordings, we cannot definitively rule out the influence of unmeasured individual differences in sleep macrostructure or quality across our sample. While our behavioral timeline was strictly controlled to target specific retention windows, future studies should incorporate physiological sleep monitoring, such as wearable devices, to clarify the precise macrostructural features of sleep that drive these memory transformations and ensure that participants followed the provided instructions regarding their sleep schedules. Importantly, despite the fact that the study was conducted virtually, results are comparable with those obtained in an in-lab study using the same design (Tompary et al., 2023), and performance was well above chance, as estimated in a previous online study using the same design (Tompary & Thompson-Schill, 2021). This suggests that participants engaged in learning despite the virtual setting.

Taken together, our results challenge the notion that sleep promotes generalization. Despite compelling models suggesting that this should be the case, empirical findings supporting this idea have been scarce. We contribute to this literature with a well-powered study that shows no effects of sleep on gist-based generalization across location memories, with some support for the counter-hypothesis that wakefulness supports generalization more than sleep does. A notable limitation of our study is that our task does not seem to be sleep-dependent. It may be argued that if memory in general does not benefit from sleep in this task, it would be a tall order to expect sleep to benefit generalization. Another potential limitation of our paradigm is that generalization is expressed before the delay, suggesting that it may have reached an effective ceiling, limiting sleep’s potential impact (however, note that inconsistent error – a marker of generalization – did, in fact, increase across sessions, irrespective of group). A final limitation is that despite the large sample size, there were still some differences between the groups that could constitute confounds for our analyses, including a slightly longer total delay between sessions and higher Inconsistent (relative to Consistent) error across both sessions for the Sleep group, and poorer novel object generalization for the Wake group at baseline. These limitations notwithstanding, our study is but one of several showing similar null effects (Lerner & Gluck, 2019). From a methodological standpoint, our study highlights the need to incorporate valid operationalizations for generalization, which ideally do not pit it against item specificity.

## Materials and methods

### Participants

A total of 534 participants were recruited and pre-screened through Prolific (Palan & Schitter, 2018), an online research platform for recruiting and managing participants. The initial pre-screening criteria required participants to: (1) be located in the Pacific Time Zone, (2) demonstrate an understanding of the study’s basic requirements by correctly answering two questions, and (3) express interest in participating in the study. Participants who failed to meet any of these criteria were excluded from the study (Figure 5). Participants were not informed of their group assignment (i.e., Sleep vs. Wake) during recruitment to avoid attrition biases.

**Figure 5:**
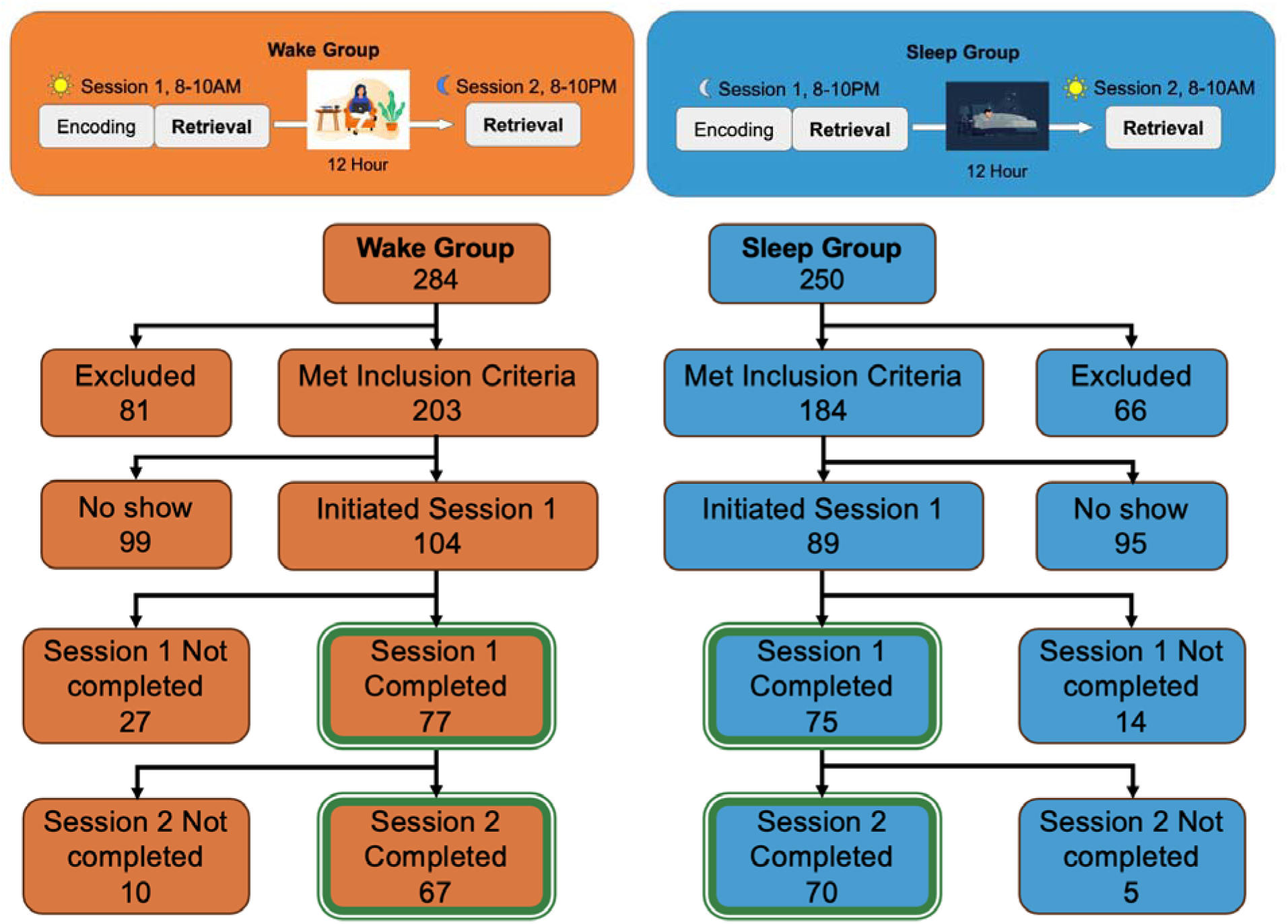
Breakdown of participant recruitment and omissions per group

Two participant groups were recruited: the Wake group (n = 284) and the Sleep group (n = 250), both completing two sessions. Participants in the Wake group were required to complete Session 1 in the morning, between 8:00 and 10:00 am, and Session 2 between 8:00 and 10:00 pm; participants in the Sleep group were asked to complete Session 1 in the evening, between 8:00 and 10:00 pm, and Session 2 the next morning between 8:00 and 10:00 am. A total of 137 participants completed both sessions of the study and were included in data analysis, comprising 67 participants in the Wake group (34 men and 33 women; mean age = 36.9 ± 11.2 SD; range 19-75 years) and 70 participants in the Sleep group (29 men, 38 women and 3 non-binary/other; mean age = 39.7 ± 12.5 SD; range 21-72 years). The groups did not differ in terms of age (*t* (135) = 1.34, *p* = 0.18) or gender ( ^2^ = 2.52, *p* = 0.28).

While a large portion of initially registered individuals did not initiate the study, the vast majority of those who did initiate it completed both sessions (Figure 5). Of those who started Session 1, 84% of participants in the Sleep group and 74% of participants in the Wake group completed that session. Of those who completed Session 1, 93% of participants in the Sleep Group and 87% of participants in the Wake Group completed Session 2.

Compensation distribution varied slightly across participants. All participants were paid 0.20 USD after completing the pre-screen survey. Participants received a total of $15 for completing both sessions of the study. For a subset of early participants (16 Sleep group participants and 14 Wake group participants), compensation was distributed so that $5 was delivered after Session 1 and $10 was delivered after Session 2. However, we then decided that the longer session - Session 1 - should be compensated at higher rates. For all subsequent participants, compensation was distributed evenly ($7.50 per session). Total compensation did not differ among participants who completed the study. Participants consented to participate in the study using an online form. The University of California, Irvine IRB approved all experimental protocols.

### Materials

The materials and study design were similar to those described in (Tompary & Thompson-Schill, 2021). The stimuli consisted of 160 color images, each measuring 100 × 100 pixels and presented on a white background. The images were evenly split into two superordinate categories: 80 animals and 80 objects. Each of these categories was further divided into four subcategories, with 20 images in each: birds, insects, sea creatures, mammals; and clothes, furniture, kitchen items, and office supplies. The typicality of the items within each category was determined using a list-ranking procedure, as described in (Tompary & Thompson-Schill, 2021).

The white background was split into two halves, with animals displayed on one side and objects on the other. Each half was further divided into four quadrants, and the four subcategories were randomly assigned to different quadrants. Across categories, all images were evenly spaced across the grid. Both the placement of the animal and object sides and the arrangement of the quadrants were randomized across participants.

Fourteen images were placed in a cluster around the center of the quadrant, while two category-typical images (e.g., Cardinal) and three category-atypical (e.g. Ostrich) images were placed outside of this cluster in a random location. To assess whether participants developed categorical knowledge, we also included a novel image placement test. Of the 20 images in each category, the most typical image was excluded from the Learning Phase in Session 1 (see below) and was only presented in the Test Phases in Session 1 and Session 2.

Stimuli were presented using customized scripts written in HTML and JavaScript. Prolific (Prolific Academic Ltd., London, UK) was used to interface with Qualtrics (Qualtrics, Provo, UT) to post pre-screen surveys and collect initial participant information. Prolific was used to interface with Pavlovia (Open Science Tools, Nottingham, UK) to post experiments, retrieve data, and pay participants.

### Procedure

The study was conducted over multiple weeks, with participants assigned to either the Wake group or the Sleep group. Each week, participants were recruited via a pre-screen survey posted on Prolific on Monday at 1:00 pm (Pacific Time). Participants who met the inclusion criteria were invited to participate in the experiment. Participants were notified of their group assignment after being accepted into the study.

For both groups, participants had until the end of Friday of the same week to complete both experimental sessions. Participants were allowed flexibility in choosing when to start Session 1, provided that it adhered to the time restrictions for their group (e.g., the Sleep group had to start between 8:00 and 10:00 pm), that Session 2 was completed ∼12 hours later, and that both sessions were completed by the end of the day Friday. The sessions were completed online, and participants were sent a reminder message if they had not started Session 1 by Thursday.

The Wake group participants were asked not to nap during the day, and the Sleep group participants were guided to have normal sleep at night. Participants were directed to use a computer / laptop to access the experiment link. All participants were directed to refrain from alcohol and tobacco use, and were asked to give undivided attention during the task. They were informed that they would not be compensated if they did not give undivided attention to the task.

After the participants accepted the Consent Form, they were directed to resize their browser window to make sure the experiment screen was fully displayed. During the Learning Phase, participants were shown an image at the bottom of the screen, and a red dot on the screen indicating the correct location of that image. Participants were asked to drag the image to the red dot, and then press continue. Trials were self-paced. There were three rounds of learning, each including 132 images equally divided between the eight categories. The order of images was randomized for each round, and participants were allowed to take a break between rounds. After the Learning Phase was completed, participants were instructed to complete a 5-min math task, which involved basic arithmetic and acted as a distraction task. After the math task, participants entered the retrieval Test Phase, where images were displayed at the bottom of the screen, and there were no red dots on the screen. Participants were asked to drag the images to the location on the screen that they remembered. Participants were not told that there would be novel images that they had not learned in Session 1. Trials were self-paced, and no feedback was provided. Because the number of images was different in the Learning and Test Phases, participants were not informed of the total number of images during learning. To provide progress feedback while maintaining consistency across phases, participants were shown only a percentage indicating task completion progress. After the Test Phase, participants filled out a questionnaire asking about basic demographic information, last night’s sleep duration and timing, participants’ degree of sleepiness, sleep habits, and whether they noticed any categories for the images presented. If they responded ‘Yes’, there additional questions were presented, inquiring about the number of categories they noticed, whether they noticed patterns, and asking that they describe these patterns. Due to a technical error, most of these questions were not required fields, and many participants neglected to fill them out. Therefore, responses to these questions were not analyzed.

Finally, participants in the Wake group were reminded not to nap during the day, and participants in the Sleep group were reminded to sleep as usual. Both groups were reminded to complete Session 2 after 12 hours.

Participants started Session 2 ∼12 hours later. In Session 2, participants completed a Test Phase, which was identical to that presented in Session 1, with the image order randomized. They then completed an additional questionnaire, asking about their last sleep bout, and whether they noticed categories / patterns. Again, data collection was inadequate, and data were not analyzed.

### Statistical analysis

All statistical analyses were conducted in MATLAB (Version R2023a, MathWorks Inc., Massachusetts). Memory accuracy was quantified for each image using the Euclidean distance between an image’s veridical location (i.e., where it was presented when encoded) and its retrieved location. Greater values of error indicate worse memory, and perfect memory would correspond to an error of 0. Generalization was quantified in several ways. We estimated the influence of category knowledge by calculating category biases (Figure 1b). First, we defined the category cluster center as the average location of the 14 Consistent images for each category. Bias was defined as the proportion of total error in the direction of an image’s category cluster (Tompary & Thompson-Schill, 2021). To compute this, we first calculated each image’s unadjusted bias by subtracting the Euclidean difference between its retrieved location and its cluster center from the Euclidean difference between its veridical location and its cluster center (Figure 1c, right). Then, we divided this unadjusted bias by the amount of error for the image. Thus, a bias score between 0 and 1 indicates that retrieval is biased toward the image’s cluster center, and a score between −1 and 0 indicates that retrieval was biased away from the cluster center. Another measure of generalization considered the placement of a typical image which was not included in the Learning Phase (i.e., a novel object). This object had no veridical location, and generalization was quantified as the distance between the retrieval location and the category cluster center (i.e., lower distances would index generalization).

For each participant and session, error was averaged across trials for all consistent, inconsistent-typical, and inconsistent-atypical images (14, 2, and 3 images, respectively). Bias was similarly averaged for all inconsistent images. Error and bias were compared between groups and sessions using mixed linear regression models, with a random effect of participant. Most of these models included group and session as fixed effects. Models examining pre-delay group differences were assessed using a single fixed effect of group. The effects of category typicality (typical vs. atypical) and spatial consistency (consistent vs. inconsistent) were analyzed using models incorporating three fixed effects (group, session, consistency/typicality). Interactions were considered for all models with two or three fixed effects.

To test for a potential tradeoff between memory specificity and generalization, we calculated for each Inconsistent image in each test whether it was placed closer to its veridical location or closer to its corresponding category center. We then compared the number of inconsistent objects placed closer to their veridical location, considering groups, sessions, and their interactions using mixed linear models, with a random effect of participants. The same was done for typical and atypical images by considering the additional fixed effect of typicality.

To test whether object placement was significantly better than chance at Session 1, we ran a permutation test: We shuffled object placements 1,000 times for each participant, calculated the mean total error for each permutation, and then compared the average error across permutations with the real error using a paired t-test. To test whether novel object placement was significantly better than chance at Session 1, we ran a permutation test: For each of 1,000 permutations, we randomly chose eight participant-indicated placements for non-novel objects. Using these positions as surrogates, we calculated the mean error across novel objects for each permutation, and then compared the average error across permutations with the real novel-object error using a paired t-test.

Results revealed differences in novel object placement in Session 1. In a post-hoc test to test whether novel object placement was affected by pre-delay differences, we ran a permutation test: We randomly eliminated 10 participants from each group 10,000 times, and compared whether a novel object distance was different between groups in Session 1 for each permutation. We then chose 10% of the permutations that had the lowest pre-delay between-group variability in distances. For these datasets, we tested whether novel object placement was affected by Session and Group with a mixed linear regression model.

## Data sharing

Data and codes will be made available upon reasonable request.

## Acknowledgments

ES is supported by the US National Institutes of Health [grant number R00-MH122663] and the US National Science Foundation [grant number 2440675]. The authors wish to thank Matthew Mak for his advice on recruitment.

## Conflict of Interest

The authors declare no conflict.

## Notes

### Competing Interest Statement

The authors have declared no competing interest.

### Summary of Updates

Revisions after reviewer comments, including mostly the Results section

